# Positional encoding in cotton-top tamarins (*Saguinus oedipus*)

**DOI:** 10.1101/186692

**Authors:** Elisabetta Versace, Jessica R. Rogge, Natalie Shelton-May, Andrea Ravignani

## Abstract

Strategies used in artificial grammar learning can shed light into the abilities of different species to extract regularities from the environment. In the *A(X)*^*n*^*B* rule, *A* and *B* items are linked but assigned to different positional categories and separated by distractor items. Open questions are how widespread is the ability to extract positional regularities from *A*(*X*)^*n*^*B* patterns, which strategies are used to encode positional regularities and whether individuals exhibit preferences for absolute or relative position encoding. We used visual arrays to investigate whether cotton-top tamarins (*Saguinus oedipus*) can learn this rule and which strategies they use. After training on a subset of exemplars, half of the tested monkeys successfully generalized to novel combinations. These tamarins discriminated between categories of tokens with different properties (*A*, *B*, *X*) and detected a positional relationship between non-adjacent items even in the presence of novel distractors. Generalization, though, was incomplete, since we observed a failure with items that during training had always been presented in reinforced arrays. The pattern of errors revealed that successful subjects used visual similarity with training stimuli to solve the task, and that tamarins extracted the relative position of *As* and *Bs* rather than their absolute position, similarly to what observed in other species. Relative position encoding appears to be the default strategy in different tasks and taxa.

## 1. Introduction

Extracting regularities is necessary to make sense of the numerous stimuli available in the environment. The relative location of different items in time and space is important in domains as different as causal reasoning (A hit B vs. B hit A), spatial navigation (A to the left of B), language (“A hit B” vs “hit B A” vs “B hit A”), action planning (“grasp A, then pierce B”). Comparing strategies and constraints in learning positional regularities across species is a way to understand cognitive species-specificities and shared abilities to process environmental regularities, including linguistic stimuli (Chen, van Rossum, & ten Cate, 2014; Fitch, 2017; Fitch & Friederici, 2012; Ghirlanda, Lind, & Enquist, 2017; Stobbe, Westphal-Fitch, Aust, & Fitch, 2012).

In the last years, attention has been posed on the implications of artificial grammar paradigms to investigate abilities relevant for language processing. In this context, the investigation of positional categories (e.g. the *A* category of items located in first position vs. the *B* category of items located in second position) has a long history. Smith (1966) exposed human subjects to four sets of letter pairs built from four classes of letters – M, N, P, and Q–, that formed MN and PQ sequences. When tested in free recall, subjects produced more intrusions of the form MQ and PN and less MP and QN intrusions than expected by chance. These results are in line with a positional encoding of the items. Some questions remained open: (i) to which extent the capacity to encode positional regularities between categories of items is widespread among different species, (ii) whether the relative or absolute position of the items is preferentially encoded.

Some studies have investigated capabilities of positional rule learning in non-human animals. Rats can use the serial/temporal order of sequentially presented elements to discriminate among them. In particular, rats can encode the sequential structure of two elements, as in A→B (Murphy et al., (2004), or three elements, as in XYX vs XXY and YXX (Murphy et al. (2008). Using long distance dependencies, Endress et al. (2010) found that violations at the edges are more salient for chimpanzees than violations within the sequence of items. These results indicate a possible specialization in processing items located at the edges. The aforementioned studies presented stimuli sequences showing tokens sequentially, thus introducing memory and attention requirements that could interfere with the performance and comparison of computational capacities across species (Fitch, 2014; Fitch, Friederici, & Hagoort, 2012; Frank & Gibson, 2008). This issue can be overcome by using visual arrays, with patterns defined by spatial relationships and all the relevant components presented at the same time. Fiser and Aslin (Fiser & Aslin, 2001, 2002, 2005) showed that after a familiarization phase, human subjects can use the frequency of occurrence of single shapes, the shape position in an array, and the arrangement of shape pairs to discriminate between familiar and unfamiliar configurations. These results suggest that computations operating on serial stimuli can be available also for visual configurations.

In the visual modality, the capacity to process rules using visual arrays has been investigated in different species (Grainger, Dufau, Montant, Ziegler, & Fagot, 2012; Murphy et al., 2008; Ravignani & Sonnweber, 2017; Rey, Perruchet, & Fagot, 2012; Scarf et al., 2016; Sonnweber, Ravignani, & Fitch, 2015; Stobbe et al., 2012; Versace, Regolin, & Vallortigara, 2006; Versace, Spierings, Caffini, ten Cate, & Vallortigara, 2017) but the processing of positional regularities has just started to be clarified. Here we use visual arrays to test the ability of a small non-human primate, the cotton-top tamarin (*Saguinus oedipus*), to encode the positional regularity *A(X)*^*n*^*B* (Chomsky, 1956). According to this grammar, *A* and *B* items are linked but assigned to different positional categories, with *As* located to the left of *Bs*. The *A* and *B* categories exhibit the positional regularities of the first and last tokens of the MN-PQ grammar introduced by Smith (1966). We used three category *A* tokens – A_1_, A_2_, A_3_ –, four category *B* tokens – B_1_, B_2_, B_3_, B_4_ –, and several *X* distractor tokens (Figure 1). The full mastery of this positional grammar requires subjects to treat familiar and unfamiliar *A(X)*^*n*^*B* combinations as grammatical and familiar or unfamiliar *B*(*X*)^*n*^*A*, *A(X)^n^A* and *B*(*X*)^*n*^*B* tokens as ungrammatical. We first trained tamarins on a subset of configurations, reinforcing choices of stimuli consistent with the *A(X)*^*n*^*B* pattern, and giving no reinforcement for choices of inconsistent stimuli. Then, we tested tamarins presenting novel items consistent and inconsistent with the rule.

**Figure 1.**
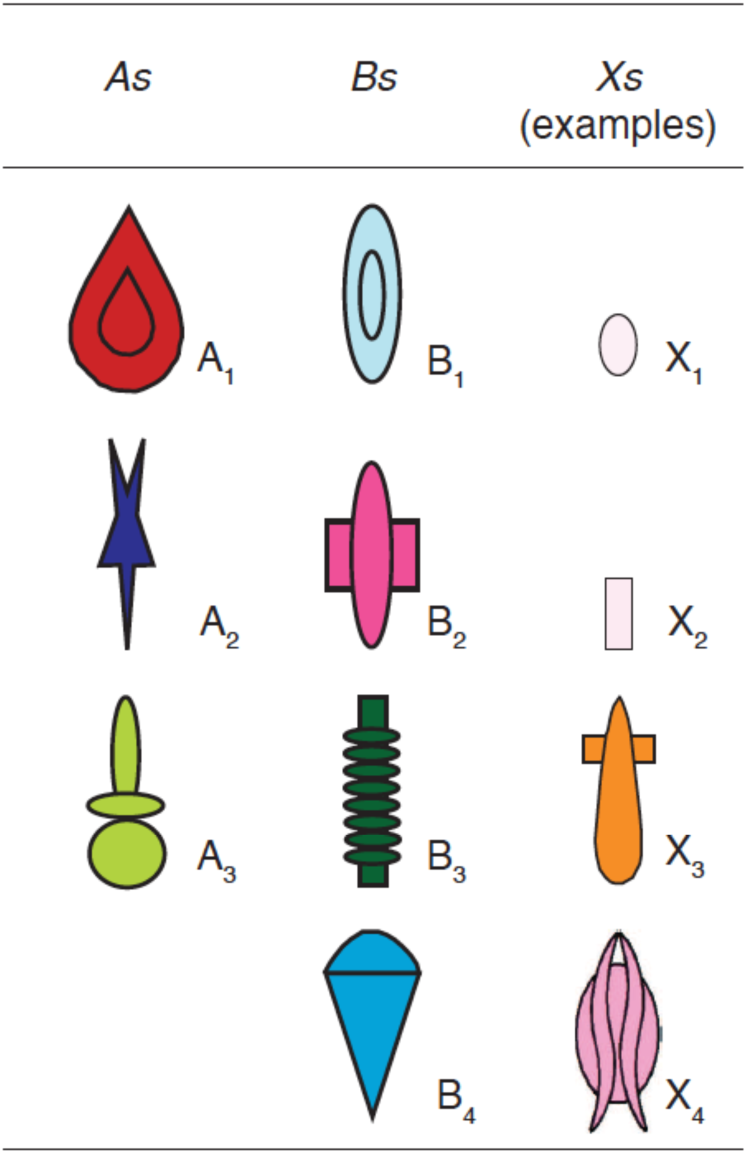
The figure shows all *A* and *B* tokens used in the experiment and four examples of *X* tokens. Arrays consistent with the rule have *A* tokens located to the left of *B* tokens. Distractor *X* tokens have no relevance in determining the consistency of an array with the grammar.

*A(X)*^*n*^*B* configurations are compatible with both absolute position rules (“*A* located on the left edge”) and relative position rules (“*A* to the left of *B*”). We used test items with distractors located at the edges (e.g. *XAXXBX* vs. *XBXXAX*) to clarify whether an individual trained on recognizing *A(X)*^*n*^*B* configurations had extracted the absolute or relative position of *As* and *Bs* with respect to the edges. Moreover, by varying the number of central distractors, we probed the relative/absolute encoding with respect to the center of the array.

## 2. General methods

The experimental schedule alternated training and test sessions. We first trained tamarins on a subset of stimuli consistent or inconsistent with the *A(X)*^*n*^*B* rule. This rule specifies the relative position of *A* and *B* tokens. We used three category *A* tokens (A_1_, A_2_, A_3_), four category *B* tokens (B_1_, B_2_, B_3_, B_4_) and a total of seventeen category *X* tokens (Figure 1).

According to the *A(X)*^*n*^*B* rule: (i) *A* tokens must be presented in left position with respect to *B* tokens; (ii) *B* tokens must be presented to the right of *A* tokens; (iii) *X* tokens can vary in number and their position is irrelevant to define the grammaticality of the configuration. When located between *As* and *Bs*, *X* tokens allow us to investigate non-adjacent relationships between *As* and *Bs*. Figure 2 shows some examples of the arrays used during the experiments, as A_2_(X_2_)^4^B_3_, A_3_(X_1_)^4^B_2_ and A_1_(X_1_)^2^B_1_ for the grammatical arrays (Figure 2a). In ungrammatical arrays (Figure 2b), the relative position of *A* and *B* tokens within the visual array was swapped, as in B_3_(X_2_)^4^A_2_ (transposition), or one token was misplaced as in B_4_(X_1_)^4^B_2_ (substitution, with two identical tokens from the same *B* category) or A_1_(X_1_)^2^A_1_ (substitution with two different tokens from the same *A* category).

**Figure 2.**
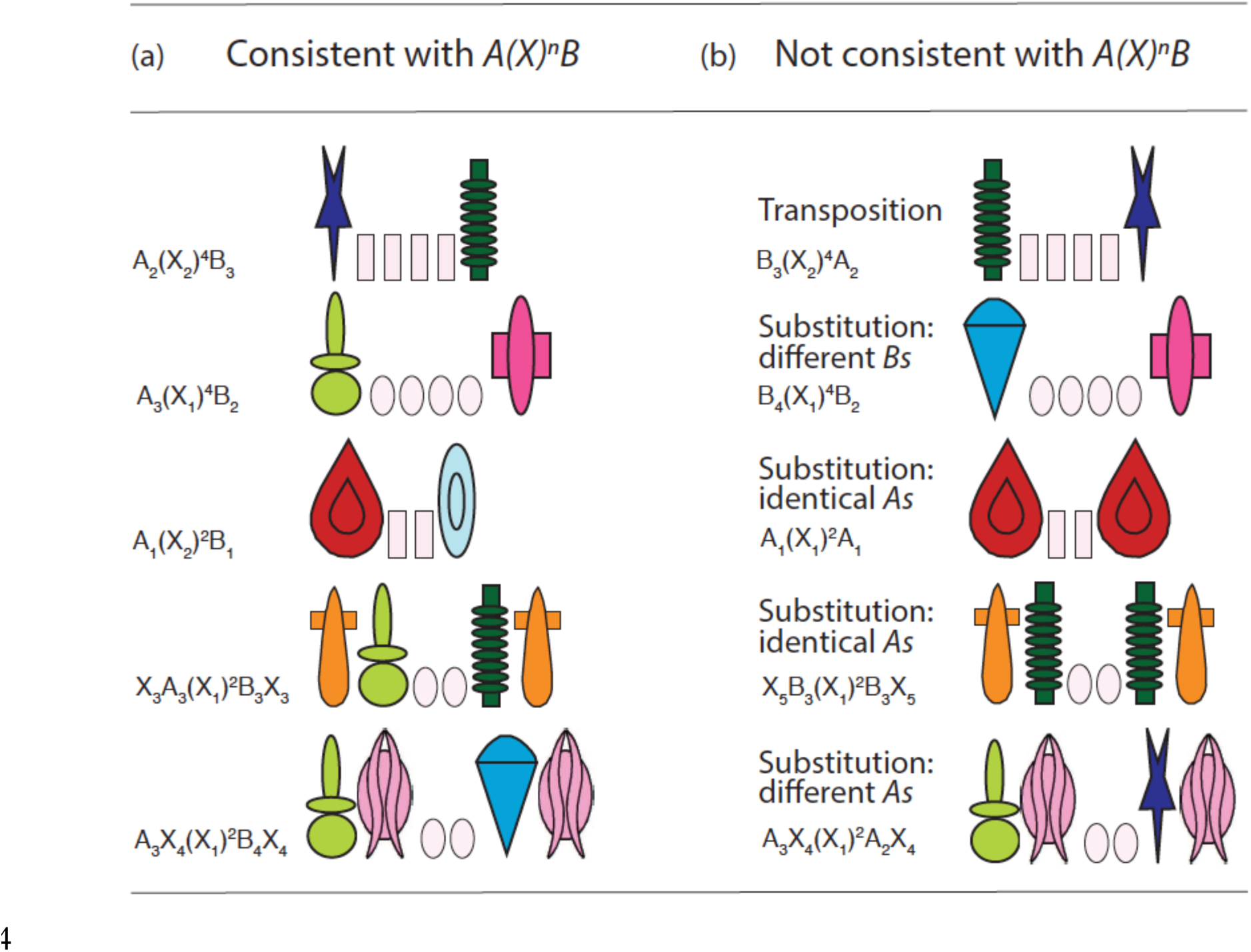
(a) Examples of arrangements consistent with the *A*(*X*)^n^*B* rule. (b) Examples of arrangements not consistent with the *A*(*X*)^n^*B* rule. In *transposition* violations, the position of *As* and *Bs* is swapped, in *substitution* violations two tokens of the same category are presented in the same array, so that only one is correctly located with respect to the other. *Identity* violations show two identical *A* or *B* tokens. Distractor tokens could be located on the edges or within the array, thus modifying the absolute position of *As* and *Bs*.

During the initial training (Training A) we presented subjects with a subset of the possible *A(X)*^*n*^*B* combinations and violations of the rule, rewarding only the choices of grammatical stimuli. In the subsequent phases, subjects were tested and trained with new combinations of the tokens used in Training A and with new *X* tokens. In particular, in Test 1 we introduced novel arrangements of *A* and *B* tokens; in Test 2 we presented new *X*s in the middle of the arrays. Finally, to assess whether each subject used a relative or absolute position strategy to solve the task, in Tests 3 and 4 we placed novel *Xs* in different locations both on the edges and within the array.

Test 1 was designed to test tamarins’ capability to extract and generalize the regularity presented during the training. Yet, succeeding in Test 1 does not clarify whether tamarins encoded the relative position of *A vs. B* tokens (“*A* to the left of *B*”) or the absolute position of *A* and *B* with respect to the edges (“*A* in the extreme left position, *B* in the extreme right position”). In fact, either the relative and absolute position encoding were consistent with the stimuli presented during the initial training. The strategy used to solve the task, more than the mere success, can inform us about representations, computational processes and biases of the subjects. Tests 2 through 4 were designed to investigate whether and how tamarins extracted a relative position or an absolute position regularity from the stimuli presented during the Training.

### 2.1. Subjects

We tested four adult cotton-top tamarins, one male (RK) and three females (RB, SH and EM), housed at Harvard University. All subjects were born in captivity and socially housed, with separate home cages for each breeding pair and their offspring. Subjects were maintained on a diet of monkey chow, fruit, seeds, and mealworms, together with free access to water. Subjects voluntarily left their home cages, lured out by a piece of raisin.

### 2.2. Apparatus

During the experimental sessions, subjects were housed in a wire mesh box (31x31x25 cm). They could access two pulling tools presented on an acrylic apparatus (40x50x6 cm) through two small holes. Each tool consisted of a pulling stem and a card covering a tray at the end of the stem. When subjects pulled one of the stems the tool advanced, the card flipped back, presenting either the food reward (a small piece of a Froot Loop© cereal) or nothing at all. Stimuli were presented on a plastic laminated sheet (11.5x7.5 cm) with different, linearly arranged shapes corresponding to the consistent or inconsistent arrays.

### 2.3. Stimuli

Tokens used to compose visual arrays belonged to three categories (*A*, *B* and *X*). Each category contained distinctive tokens, as shown in Figure 1. Tokens were arranged in visual arrays, printed on cards and located on the apparatus. The tokens used and their position within the array determined whether the stimulus printed on each card was consistent or inconsistent with the target rule.

Consistent arrays followed the *A(X)*^*n*^*B* rule, for example A_1_X_1_X_1_B_2_. In arrays not consistent with the target rule, the position of *A* and *B* tokens was swapped, as in B_2_X_1_X_1_A_1_, or either *A* or *B* tokens were not used. *X* tokens, irrelevant in determining the consistency of the stimuli with the target rule, could vary in size and number, extending or reducing the distance of the dependency between other tokens.

Possible arrangements of the *A* and *B* tokens are shown in Table 1: 12 patterns are consistent with the target rule *A(X)*^*n*^*B* and 37 are not consistent with it. Arrangements not consistent with the rule include transpositions, in which *Bs* are located to the left of *As*, and substitutions, in which two tokens of the same A (or B) category are present. During the initial training, we used only 6 of the grammatical arrangements and 7 of the ungrammatical arrangements, 2 or 4 X_1_ and X_2_ tokens, saving the other configurations of stimuli for the tests. The 13 patterns employed in the training are shaded grey in Table 1, and shaded in green and red in the electronic version.

**Table 1.** Possible arrangements of *As* and *Bs*. Only shaded patterns were used during the training. Consistent patterns follow the *A*(*X*)^n^*B* rule, inconsistent pattern swap the position of *As* and *B*s (transpositions) or show two different or identical tokens of the same *A* or *B* category (substitutions).

| Consistent<br>with $A(X)^nB$ | Transposition | Substitution:<br>Different $A$s | Substitution:<br>Identical $A$s | Substitution:<br>Different $B$s | Substitution:<br>Identical $B$s |
| --- | --- | --- | --- | --- | --- |
| $A_1(X)^nB_1$ | $B_1(X)^nA_1$ | $A_1(X)^nA_2$ | $A_1(X)^nA_1$ | $B_1(X)^nB_2$ | $B_1(X)^nB_1$ |
| $A_1(X)^nB_2$ | $B_1(X)^nA_2$ | $A_1(X)^nA_3$ | $A_2(X)^nA_2$ | $B_1(X)^nB_3$ | $B_2(X)^nB_2$ |
| $A_1(X)^nB_3$ | $B_1(X)^nA_3$ | $A_2(X)^nA_1$ | $A_3(X)^nA_3$ | $B_1(X)^nB_4$ | $B_3(X)^nB_3$ |
| $A_1(X)^nB_4$ | $B_2(X)^nA_1$ | $A_2(X)^nA_3$ | | $B_2(X)^nB_1$ | $B_4(X)^nB_4$ |
| $A_2(X)^nB_1$ | $B_2(X)^nA_2$ | $A_3(X)^nA_1$ | | $B_2(X)^nB_3$ | |
| $A_2(X)^nB_2$ | $B_2(X)^nA_3$ | $A_3(X)^nA_2$ | | $B_2(X)^nB_4$ | |
| $A_2(X)^nB_3$ | $B_3(X)^nA_1$ | | | $B_3(X)^nB_1$ | |
| $A_2(X)^nB_4$ | $B_3(X)^nA_2$ | | | $B_3(X)^nB_2$ | |
| $A_3(X)^nB_1$ | $B_3(X)^nA_3$ | | | $B_3(X)^nB_4$ | |
| $A_3(X)^nB_2$ | $B_4(X)^nA_1$ | | | | |
| $A_3(X)^nB_3$ | $B_4(X)^nA_2$ | | | | |
| $A_3(X)^nB_4$ | $B_4(X)^nA_3$ | | | | |

Test 1 featured test stimuli consisting of novel arrangements of the tokens previously used during the training (unshaded stimuli in Table 1). In Test 2 we introduced four novel *X* tokens in the central position (we used for example stimuli such as A_1_(X_3_)^n^B_2_ and A_2_(X_4_)^n^A_2_). In Test 3 and Test 4 we introduced new *X* tokens in novel positions either internally (as A_3_X_6_(X_1_)^2^B_4_X_7_ and A_3_ X_6_(X_1_)^2^X_7_A_2_ in Test 3) or at the edges of the arrays (as X_6_A_3_(X_1_)^2^B_3_X_7_ and X_6_B_3_(X_1_)^2^A_1_X_7_ in Test 4, see Figure 2). *Xs* were irrelevant in determining the rule-consistency of the stimuli and were used to evaluate the information encoded by tamarins during the training.

### 2.4. Procedure

Before starting the training on the *A(X)*^*n*^*B* rule, each subject was familiarized with the apparatus and the experimental procedure. Both training and test sessions involved the following procedure. The target subject was lured out of its home cage with a piece of food into a transport box and then moved individually to the experimental room for an experimental session. Prior to a trial, and out of view from the subject, the experimenter prepared the appropriate stimuli and reward. The overall sequence of different stimulus pairings, along with the right or left position of each card, was randomized and counterbalanced across trials and within sessions. For each session, consistent cards were equally distributed between the right and left sides of the apparatus and no more than two consistent cards were presented on the same side consecutively.

The apparatus was presented for three seconds in a position out of reach from the subject at a distance of approximately 40 cm, and then subsequently pushed towards the subject. In cases where the subject did not look at the setup within four seconds, the experimenter drew the subject’s attention to the tray by pointing to the midpoint between the cards, and then moved the tray forward. The subject was only allowed to pull one of the two tools; following its first selection, the alternative tool was retracted, out of reach. When the subject pulled the tool with the consistent card, a food reward was immediately available. If the subject pulled the tool with the inconsistent card, the experimenter revealed the hidden food under the consistent card. In both training and testing sessions, correct choices were rewarded in the same way. Soon after the subject completed its choice, the apparatus was removed and the experimenter prepared a new trial out of view. If the subject did not make any choice within 8 seconds, the apparatus was removed and the trial aborted.

Each training session consisted of 2 to 6 warm-up trials followed by 12 training trials. Warm-up trials continued until 2 consecutive correct choices were made, at which point the experimenter proceeded the session. If more than 6 warm-up trials were necessary, the session was aborted. Warm-up trials consisted of consistent stimuli that the subject had successfully discriminated in previous conditions, and were designed to make sure that on each session the subject was attentive and motivated; as such, if a subject were attending to the material and motivated to pull the tool, it should succeed on the warm-up trials.

Each test session consisted of 2-6 warm-up trials followed by 12 test trials interspersed with 4 trials with stimuli already presented during the previous training. When responsive, each subject ran two experimental sessions per day. To guarantee an appropriate level of motivation, the inter-session interval within a day was at least three hours. The difference between training and test sessions consisted in the novel material presented during test sessions.

Monkeys’ responses were coded in terms of which array (consistent or inconsistent) was selected on each trial. To move from the initial training stage (Training 1) to the tests, we required subjects to reach a criterion of 40/48 correct trials or better. This corresponds to the cut-off value for a binomial test with alpha=0.05, consisting of 10 out of 12 correct trials over 4 consecutive sessions or better.

To determine whether subjects could discriminate between novel consistent and inconsistent stimuli, showing generalization, we analyzed (a) the scores of the first 48 trials (4 sessions) and (b) the scores of the first 96 test trials (8 sessions). We ran the analysis on 8 sessions to increase the number of trials and investigate the responses to specific violations.

### 2.5 Experimental schedule

The experimental schedule went through the following stages, summarized in Table 2: Training A, Test 1, Training B, Test 2, Test 3, Test 4.

**Table 2.** Experimental schedule, composed by trainings and test stages. Shaded rows indicate the tests.

| Stage | Stimuli |
| --- | --- |
| Training A | A subset of 13 out of 49 possible arrangements |
| Test 1 | Novel arrangements |
| Training B | Stimuli presented in the previous two stages |
| Test 2 | Novel $Xs$ in the centre |
| Test 3 | Novel $Xs$ on the edges |
| Training C | Same stimuli presented in Training B |
| Test 4 | Novel $Xs$ between $As$, $Bs$ and $Xs$ |

Training A lasted until the subject reached the criterion of at least 40/48 correct responses in four consecutive sessions. During Training A, we progressively added the experimental stimuli. In this stage, subjects were presented with all the *A* and *B* tokens but only a subset of all the possible combinations of them (see Table 1).

Test 1 explored the extent to which tamarins generalized the distinction between consistent and inconsistent stimuli to new spatial arrangements of the tokens experienced during the training. We hypothesized that if tamarins had encoded the positional regularity of *A* and *B* tokens (“*A* to the left of *B*”) they should, in the absence of further training, choose consistent combinations (i.e. *A(X)*^*n*^*B*) more often than inconsistent combinations.

Test 2 was identical to Test 1, with the exception that experimental trials were composed with four novel *Xs*. Test 2 explored whether tamarins could generalize to novel *Xs* located in the center of the arrays.

Subjects responding above chance to Test 1 and Test 2 moved on to Training B, in which we presented a total of 36 sessions to each subject, using the same stimuli presented in Test 1. Test 3 was identical to Test 1, except that new *Xs* were located at the edges of the arrays, so that stimuli followed patterns similar to *XA(X)*^*n*^*BX* or *XB(X)*^*n*^*AX*. Up to this stage, *A* and *B* tokens always occupied the edges of the sequence. Two alternative hypotheses could account for tamarins’ success until Test 2. Tamarins could have learned, instead of the relative position of *A* and *B* tokens, their absolute position with respect to the edges. In that case, they would not have been able to generalize to arrays not containing *A* or *B* tokens at the edges, as presented in Test 3. As an alternative hypothesis, if tamarins had learned that *A* must be on the left with respect to *B*, their performance should not have been affected by the insertion of novel tokens at the edges. Subjects responding above chance in Test 3 proceeded to Training C (with the same stimuli used in training B) and then to Test 4.

Test 4 was identical to Test 1, except that at least one *X* token was located between *As* or *Bs* and the central *Xs*, so that stimuli followed patterns of the form *XA(X)*^*n*^*XB* or *BX(X)*^*n*^*AX*. If tamarins learned that the position of *A* and *B* with respect to *X* tokens were irrelevant, or that the symmetry of *A* and *B* with respect to the center of the array were irrelevant, than their performance should not be affected by the insertion of novel *Xs* adjacent to the central tokens.

## 3. Results and discussion

### 3.1. Training A

Training A lasted until the subject reached the criterion of at least 40/48 correct responses in four consecutive sessions. As expected, subjects varied in terms of the number of training sessions required to reach criterion: RK required 278 sessions over 149 days of training (Mean=1.86 sessions/day), RB required 265 sessions over 209 days of training (Mean=1.27 sessions/day), EM required 234 sessions over 175 days of training (Mean=1.34 sessions/day), SH required 487 sessions over 271 days of training (Mean=1.79 sessions/day). For SH we interrupted the training when she gave birth and then restarted the training for 261 sessions over 143 days (Mean=1.81 sessions per day).

### 3.2. Test 1: Novel arrangements

Figure 3 shows each subject’s performance at the end of Training A (black line), and in Test 1 (red line) as percentage of accuracy (number of correct choices/total number of trials*100). We calculated the number of correct choices for each subject in the first four and eight test sessions (48 and 96 trials respectively) of Test 1 and tested whether the number of correct choices was significantly different from chance with a two-tailed binomial test.

**Figure 3.**
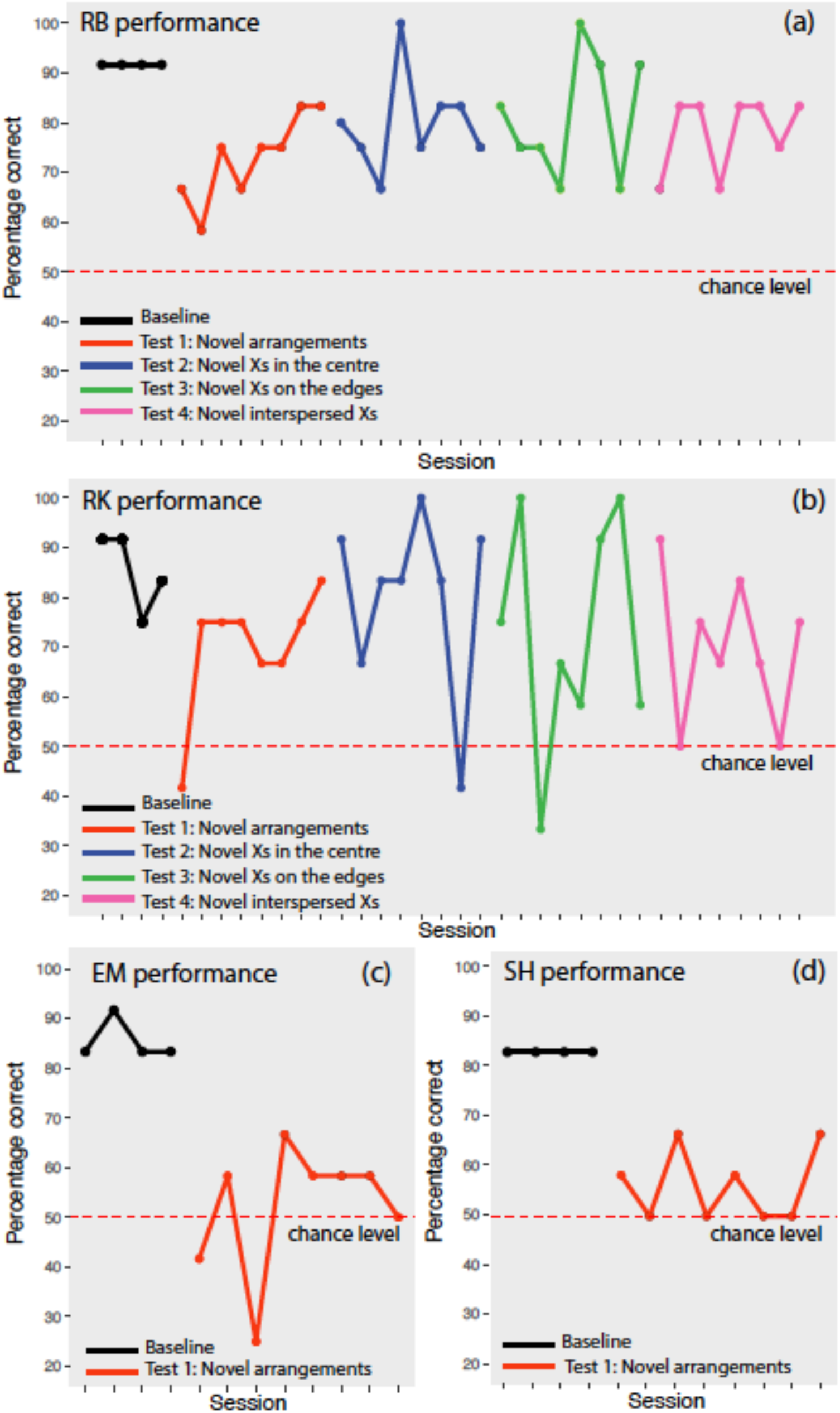
Each panel shows the performance of a subject: (a) for RB, (b) for RK, (c) for EM, (d) for SH. The baseline criterion (black line) and the first eight sessions of each test are visualized: in Test 1 (red line) subjects were probed with novel arrangements of *As* and *Bs*, in Test 2 (blue line) with novel central *Xs*, in Test 3 (green line) with novel *Xs* on the edges, in Test 4 (pink line) with novel *Xs* within the array.

In the first four sessions two subjects – RK and RB – performed significantly better than chance (RK: 32/48 correct choices, 67%, p=0.029; RB: 32/48 correct choices, 67%, p=0.029), while two subjects – EM and SH – did not (EM: 23/48 correct choices, 48%, p=0.885; SH: 27/48 correct choices, 56%, p=0.471)). Similarly, in the first eight sessions only two subjects performed significantly better than chance: RK: 67/96 correct choices, 70%, p<0.001; RB: 70/96 correct choices, 73%, p<0.001; EM: 50/96 correct choices, 52%, p=0.76; and SH: 54/96 correct choices, 56%, p=0.26.

These results license the conclusion that RK and RB (but not the other two subjects, EM and SH) were able to use the experience gained during Training A to successfully distinguish between novel consistent and inconsistent stimuli. Thus, at least two cotton-top tamarins could learn a positional rule as *A(X)*^*n*^*B* and generalize this regularity to novel arrangements. The positive performance of RK and RB is noteworthy considering that, during the training, subjects had previous experience with only a small set of stimuli (n=13 token combinations), which then increased to a set of 24 novel exemplars presented in Test 1, and that before Test 1 tamarins had never encountered substitutions with different tokens of the same category. We ran further probe tests to investigate the encoding of the regularity in RB and RK.

### 3.2. Training B and Test 2: Novel *Xs* in the center

Before moving to Test 2, subjects that succeeded in Test 1 were trained with 36 sessions identical to those presented in Test 1 (Training B). The success rate over the 36 sessions for each subject was 328/432 correct choices, 76% for RK, and 315/420 correct choices, 75% for RB.

In Test 2, we investigated whether RK and RB were able to generalize to new *X* tokens located between *A* and *B*, in the center of the arrays. The results of the first eight sessions are presented as the blue line in Figure 3. Both monkeys performed significantly above chance (two-tailed binomial tests): RK made 77/96 correct choices, 80%, p<0.001, and RB made 75/94 correct choices (two trials were aborted because the subject was not responsive), 80%, p<0.001. The same outcomes were observed in the first four sessions: RK made 39/48 correct choices, 81%, p<0.001; RB made 37/48 correct choices, 77%, p<0.001). Hence the performance of RK and RB was not disrupted by change in *X* tokens in the center of the arrays, suggesting that these monkeys were not using the absolute position of the tokens to make their choices, and that the distractor *X* tokens in central position did not affect their choices.

### 3.3. Test 3 and Test 4: Novel *Xs* on the edges and between *As*, *Bs* and *Xs*

In Test 3 (Figure 3, blue line) novel *Xs* were added on the edges of the arrays, thus changing the absolute position of *As* and *Bs* with respect to the edges. In the first 8 sessions (96 trials) RK scored 70/96 correct choices, 73%, p<0.001 and RB scored 78/96 correct choices, 81%, p<0.001. Similarly, considering the first four sessions, RK made 33/48 correct choices, 69%, p= 0.013, and RB made 36/48 correct choices, 75%, p<0.001). Hence, tamarins responded above chance level even when the absolute position of *As* and *Bs* was changed with respect to the edges.

In Test 4 (Figure 3, green line) at least one *X* was located between *A* or *B* and the central *Xs*. In the first eight sessions of Test 4, RK made 67/96 correct choices, 70%, p<0.001; RB made 75/96 correct choices, 78%, p<0.001. In the first four sessions, RK made 34/48, 71%, p=0.006; RB made 36/48 correct choices, 75%, p<0.001). Hence, the performance of RB and RK was not disrupted by the insertion of novel *Xs* that changed the absolute position of *As* and *Bs* with respect to the center of the array and that made the stimuli asymmetrical. Both monkeys appeared to use a relative position strategy to solve the task.

### 3.4 Learning during the test and differences between test stages

In the first sessions of the tests, monkeys did not perform significantly worse than in subsequent test sessions (chi-squared test, RB: chi-squared=0.112, p=0.737; RK: chi-squared=0.017, p=0.896). Moreover, there was no significant correlation between percentages of correct responses and temporal sequence of test sessions (Spearman’s correlation, RB: t_30_=1.56, p=0.129; RK: t_30_=-0.212, p=0.834). This evidence suggests that the eight test sessions are representative of the generalization performance.

To test whether the performance of RK and RB differed across test stages, we used a chi-squared test comparing correct vs. incorrect choices in the four tests. RB and RK did not make significantly more incorrect choices across the four different test stages (RB: chi-squared_3_=2.344, p=0.504; RK: chi-squared_3_=3.457, p=0.326), see Figure 4.

**Figure 4.**
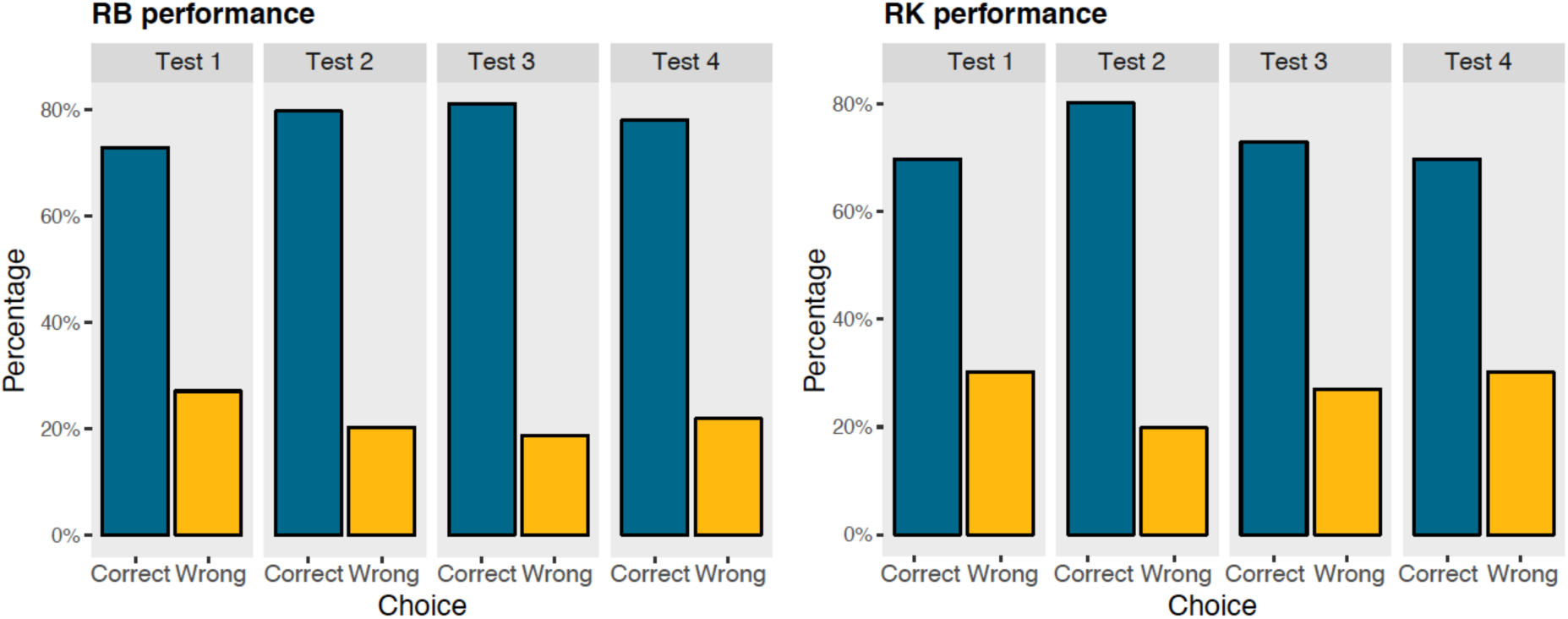
Overall performance (percentage of correct and wrong choices) by test stage for RB (left panel) and RK (right panel).

### 3.5 Analyses of responses to inconsistent stimuli

We ran further analyses to investigate the individual strategies used by the subjects that succeeded in Test 1 and went through the other test stages, by looking at the pattern of responses to arrangements which were inconsistent with the *A(X)*^*n*^*B* grammar in the first eight sessions of each test stage.

To investigate the presence of difficulties or enhanced performance in the presence of specific tokens we analyzed the responses to each *A* (Figure 4) and *B* token (Figure 5) presented in ungrammatical stimuli. As showed in Figure 4, both RB and RK performed significantly worse with inconsistent arrangements that contained A_3_ (RB: chi-squared=13.737, p<0.001; RK: chi-squared=14.462, p<0.001) than with other tokens. During training, we had not used the A_3_ token in any unrewarded arrangements (see Table 1) to test for generalization to items presented in novel positions. Hence the specific failure with A_3_ exhibited by both monkeys suggests an incomplete generalization of the positional A(X)^n^B rule that can be due to the selective inexperience with A_3_ as unrewarded token.

**Figure 5.**
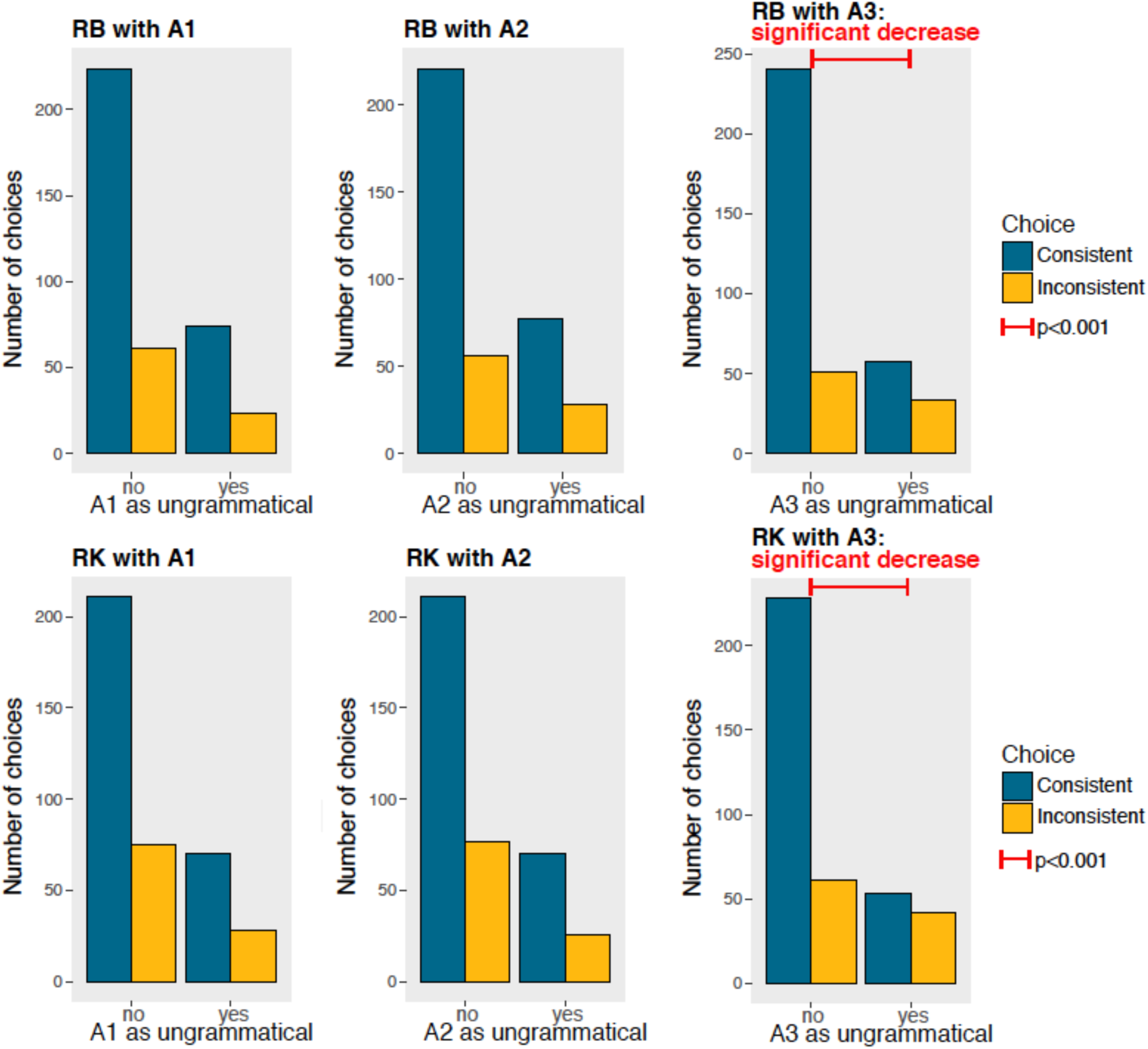
Each chart shows the performance of a subject (RB on the top panels, RK on the bottom panels) when each *A* token was present or absent in the ungrammatical arrays. Blue bars indicate correct choices, consistent with the grammar, yellow bars indicate wrong choices, inconsistent with the grammar. Both RB and RK performances were significantly lower when A_3_ tokens were presented in ungrammatical arrangements, suggesting a similar encoding of the regularity.

We did not find learning difficulties with any other token, but the performance of each monkey was enhanced in the presence of some tokens, although these effects were not consistent across subjects. These are the results with *A* tokens (Figure 5) for RB: A_1_ chi-squared=0.110, p=0.740; A_2_: chi-squared=1.490, p=0.222; and for RK: A_1_ chi-squared=0.110, p=0.749; A_2_ chi-squared=0, p=1. These are the results for *B* tokens (Figure 6) for RB: B_1_ chi-squared=4.938, p=0.026 with a significant performance enhancement; B_2_ chi-squared=1.276, p=0.259; B_3_ chi-squared=3.595, p=0.058 with a trend for performance enhancement; B_4_ chi-squared=1.292, p=0.255, and for RK: B_1_ chi-squared<0.001, p=1; B_2_: chi-squared=0.0008, p=0.977; B_3_ chi-squared=0.358, p=0.550; B_4_ chi-squared=3.235, p=0.071 with a trend for performance enhancement.

**Figure 6.**
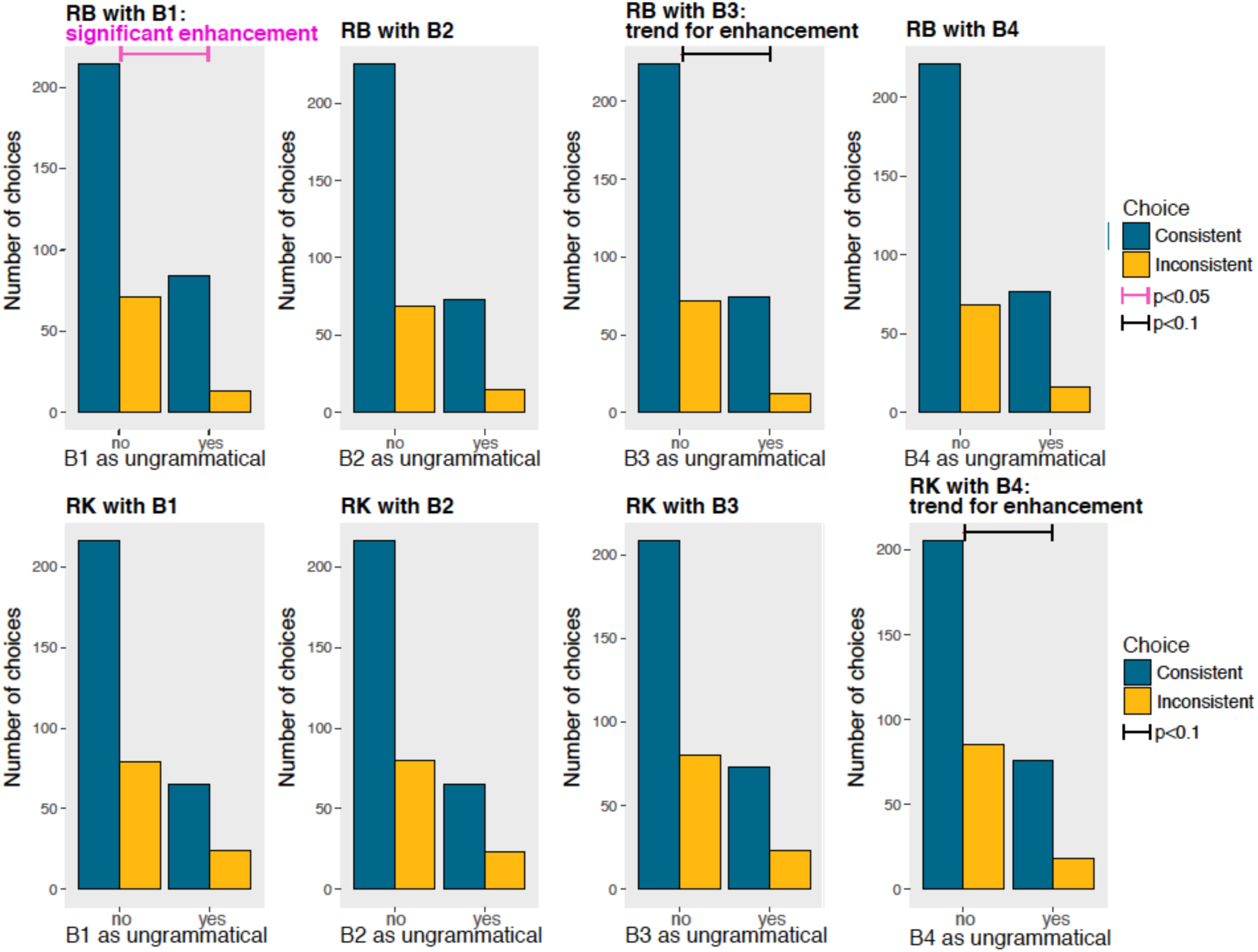
Each chart shows the performance of a subject (RB on the top panels, RK on the bottom panels) when each *B* token was present or absent in the ungrammatical arrays. Blue bars indicate correct choices, consistent with the grammar, yellow bars indicate wrong choices, inconsistent with the grammar. Both RB and RK performances were significantly lower when A_3_ tokens were presented in ungrammatical arrangements, suggesting a similar encoding of the regularity.

We analyzed the effect of different arrangement types on violations: presence/absence of transpositions (swapped position of *As* and *Bs*), presence/absence of identity violations (two identical *A* or *B* tokens), presence/absence of violations located on the edges or on the center (Figure 7).

**Figure 7.**
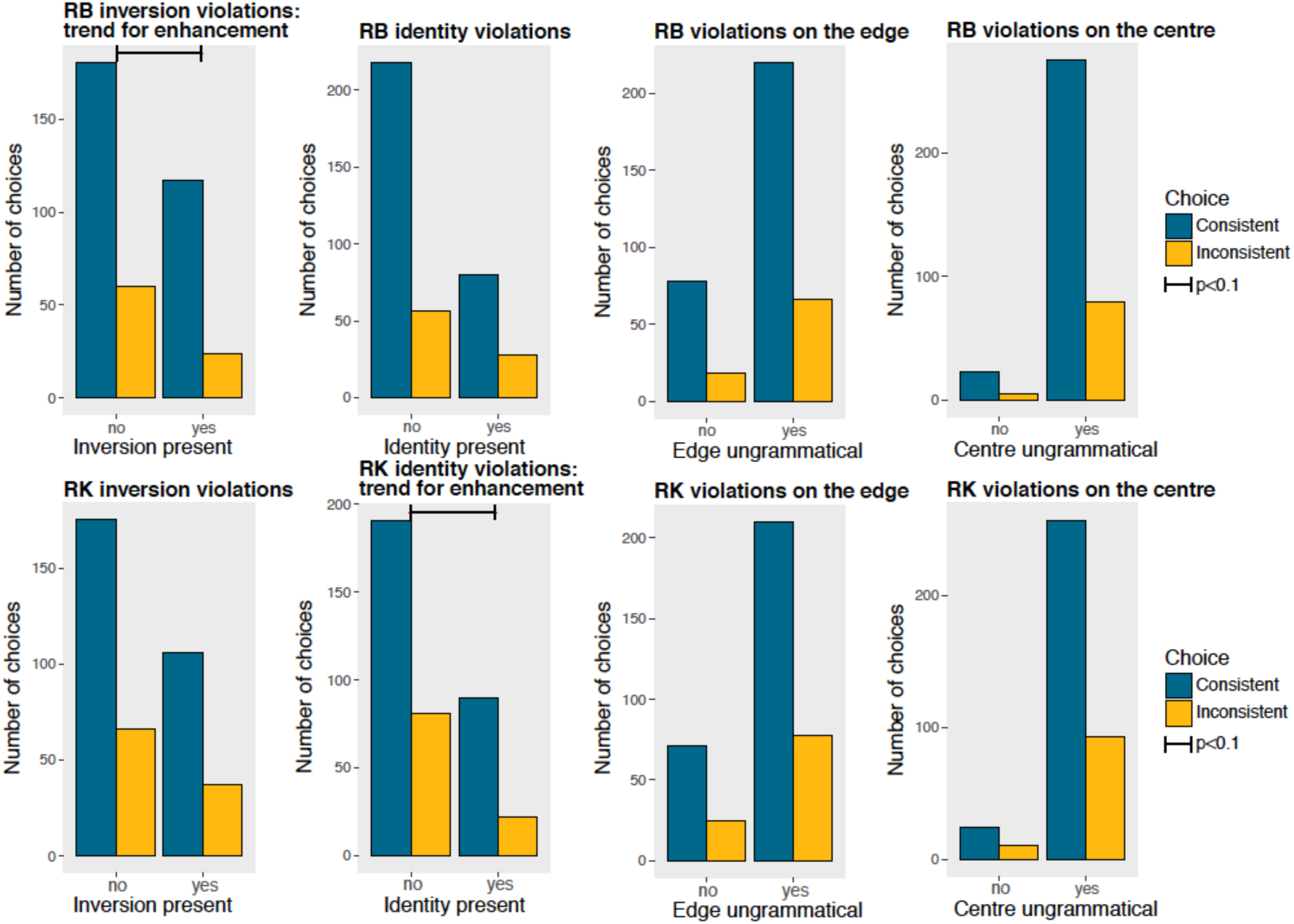
Each chart shows the performance of a subject presented with different violations: presence/absence of transpositions (swapped position of *As* and *Bs*), presence/absence of identity violations (two identical *A* or *B* tokens), presence/absence of violations located on the edges or on the center.

If monkeys were taking into account the relative position of the tokens, we would have expected a better performance with *transpositions*, in which the *A* and *B* token is swapped, thus producing a double violation, compared to *substitutions*, in which only one token is located in the incorrect position. On the contrary, we could have observed a pattern similar to the transposed-letter effect observed in human literates (Dunabeitia, Orihuela, & Carreiras, 2014; Grainger, 2008; Perea & Lupker, 2004) and pigeons (Scarf et al., 2016), in which ungrammatical letter strings (non-words) obtained by transposing adjacent letters in a grammatical letter string (word) induce misclassifications. Comparing the performance of transpositions vs. substitutions, we did not observe any significant enhancement or decrease in performance with transpositions compared to substitutions, although RB had a trend for enhancement with transpositions (RB chi-squared=2.773, p=0.096 with a trend for enhancement; RK chi-squared=0.042, p=0.838). Hence, differently from pigeons (Scarf et al., 2016) tamarins’ performance was not negatively affected by transposition, and potentially slightly enhanced.

We tested whether rejecting arrangements with substitutions with two identical tokens was easier than rejecting other violations of the grammar, and this was not the case for RB (chi-squared=1.059, p=0.303), while there was a trend for enhancement in RK (chi-squared=2.653, p=0.056). Violations on the edges (RB chi-squared=0.552, p=0.457; RK chi-squared=0.004, p=0.947) and on the center (RB chi-squared=0.097, p=0.756; RK chi-squared=0.014, p=0.878) were not significantly different than other violations. Overall, the lack of enhanced performance with any class of violation suggests that monkeys used a strategy based on configurational encoding, rather than an encoding based on the analysis of the position of each token, to respond to novel stimuli.

## 4. General discussion

Detecting in which position elements occur relative to one another is important in many domains, such as language processing and animal communication (e.g. ten Cate & Okanoya, 2012), orthographic processing (Dunabeitia et al., 2014; Scarf et al., 2016; Ziegler et al., 2013), spatial navigation and foraging (Rugani, Kelly, Szelest, Regolin, & Vallortigara, 2010; Vallortigara & Zanforlin, 1986), action planning (Raby, Alexis, & Clayton, 2007), among others. It is not clear, though, which representations and computational mechanisms different species use to encode positional relationships. In the present work, we investigated how the positional regularity *A(X)*^*n*^*B* is processed in the New World monkeys cotton-top tamarins (*Saguinus oedipus*). Only stimuli in which tokens of the category *A* are located to the left of tokens of the category *B* are consistent with this rule, while other tokens are not relevant in determining the grammaticality of stimuli. To reduce the attention and memory load required to solve the task and focus on computational capabilities and generalization, we tested monkeys using visual arrays.

Monkeys were able to learn a positional regularity between non-adjacent items such as *A(X)*^*n*^*B*. In fact, after being trained on a subset of stimuli, two out of four subjects were able to immediately generalize to novel arrangements the distinction between items consistent and not consistent with the target rule. The limited evidence provided during the training was sufficient for monkeys to tell apart the positional role of *A*, *B* and distractor *X* tokens. In another work, Grainger et al. (2012) trained baboons to categorize novel visual arrays of stimuli composed by letters arranged according to a statistical positional regularity. Monkeys were able to acquire adjacent dependencies over letter bigrams within the arrays and coded the word vs. non-word stimuli as sets of letter identities arranged in a specific order. A subsequent experiment with baboons (Ziegler et al., 2013) showed that monkeys, similarly to human readers, exhibit the transposed–letter effect (see Grainger, 2008), so that their performance was lowered by letter transpositions. In this work, though, it was not clear whether monkeys relied on an absolute or on a relative position strategy. A significant decrease in performance with transpositions mediated by relative-position encoding has been noticed in human literates (Dunabeitia et al., 2014; Grainger, 2008) and in pigeons trained to orthographical discriminations (Scarf et al., 2016). We ran several tests to clarify whether tamarins had encoded the absolute position or the relative position of *As* and *Bs* (Versace, 2008). In our experiments, the absolute position of a token within an array, such as “A_1_ must be located as first token on the left part of the array”, is defined independently from the identity of other tokens. On the contrary, its relative position, such as “A_1_ must be located to the left of B_1_, B_2_, B_3_ or B_4_”, depends on the specific identity and position of other tokens. These two alternative strategies of encoding can be probed changing the absolute position of *As* and *Bs* with respect to the edges and the center of the array by inserting novel *Xs* in different positions within the visual arrays. If during the training tamarins had encoded the absolute and not the relative position of *As* and *Bs*, they were expected to fail when the absolute position of *As* and *Bs* was changed. On the contrary, tamarins’ performance was not disrupted when novel *Xs* were added in the center of the arrays and when the absolute position of *As* and *Bs* was changed with respect to the edges and the center of the array. We can hence conclude that tamarins did not rely on the mere absolute position of *As* and *Bs*. The fact that tamarins’ performance was not compromised by transpositions suggests that they might have used a visual similarity strategy to solve the task. This conclusion is supported by the fact that both successful subjects had a significantly lower performance with ungrammatical stimuli that contained a token – A_3_ – never presented in unrewarded stimuli during the training.

Overall, we documented a preferential encoding of the relative position. Preferential encoding of relative vs. absolute position has not been documented only in primates and pigeons. Relative rather than absolute encoding in spatial positions had been previously shown in chicks of the domestic fowl during foraging (Vallortigara & Zanforlin, 1986). In a series of experiments, chicks were trained to discriminate between two boxes according to either their relative position to each other or their absolute position (the position with respect to the geometry of the cage or other features of the environment). When the boxes were located close to each other, learning on the basis of the relative position was faster than learning on the basis of absolute position. The advantage of relative position was reduced only when the boxes were located further apart. Similarly, our results obtained with closely located tokens show a preference for relative encoding of visual stimuli presented in simultaneous configurations. It seems that encoding of relative rather than absolute position is the default strategy observed across different species and taxa, irrespectively of the possess of language. Further studies should investigate this phenomenon and clarify its spread, neurobiological and evolutionary basis.

## Acknowledgements

All research was approved by the Animal Care and Use committee at Harvard University. Funds for this research were provided to E.V. by the Harvard Mind Brain and Behavior (MBB) Graduate Student Award. We thank the members of the Cognitive Evolution Laboratory (Harvard University) for help in data collection and comments on the data.

